# Development and application of a computable genotype model in the GA4GH Variation Representation Specification

**DOI:** 10.1101/2022.09.06.506817

**Authors:** Wesley Goar, Lawrence Babb, Srikar Chamala, Melissa Cline, Robert R. Freimuth, Reece K. Hart, Kori Kuzma, Jennifer Lee, Tristan Nelson, Andreas Prlić, Kevin Riehle, Anastasia Smith, Kathryn Stahl, Andrew D. Yates, Heidi L. Rehm, Alex H. Wagner

**Affiliations:** Institute for Genomic Medicine, Nationwide Children’s Hospital, Columbus, OH; Broad Institute of MIT and Harvard, Cambridge, MA; Department of Pathology and Laboratory Medicine, Children’s Hospital Los Angeles, Los Angeles,CA; UC Santa Cruz Genomics Institute, Santa Cruz, CA; Department of Artificial Intelligence & Informatics, Center for Individualized Medicine, Mayo Clinic; MyOme, Inc, San Carlos, CA; Sequencing.com, Los Angeles, CA; Geisinger, Danville, PA; Invitae, San Francisco, CA; Baylor College of Medicine, Houston, TX; European Molecular Biology Laboratory, European Bioinformatics Institute, Wellcome Genome Campus, Cambridge, UK; Massachusetts General Hospital, Boston, MA; Department of Pediatrics & Biomedical Informatics, The Ohio State University College of Medicine, Columbus, OH

**Keywords:** Genomics, GA4GH, VRS, Genotype, Haplotype, Allele, HGVS, VCF

## Abstract

As the diversity of genomic variation data increases with our growing understanding of the role of variation in health and disease, it is critical to develop standards for precise inter-system exchange of these data for research and clinical applications. The Global Alliance for Genomics and Health (GA4GH) Variation Representation Specification (VRS) meets this need through a technical terminology and information model for disambiguating and concisely representing variation concepts. Here we discuss the recent Genotype model in VRS, which may be used to represent the allelic composition of a genetic locus. We demonstrate the use of the Genotype model and the constituent Haplotype model for the precise and interoperable representation of pharmacogenomic diplotypes, HGVS variants, and VCF records using VRS and discuss how this can be leveraged to enable interoperable exchange and search operations between assayed variation and genomic knowledgebases.

## 1. Introduction

Representation of genomic variation as recorded in genomic data systems is highly varied and complex, involving the computable formalization of imprecise concepts with imprecise definitions for data exchange between systems. Several well-known formats and tools have been developed for exchanging some common forms of variation, including the Variant Call Format (VCF)^1^, the Human Genome Variation Society (HGVS) variant nomenclature^2^, the NCBI Sequence-Position-Deletion-Insertion (SPDI) data model^3^ and the ClinGen Allele Registry web service^4^, among others^5–8^. Despite this, these common fit-for-purpose variation models use unaligned terminologies, conventions, and assumptions that make it challenging to losslessly convert information between formats. More pressingly, these formats are difficult to extend to domain-specific requirements for variation representation across different communities, promoting further division of terms, information models, and exchange formats for genomic variation^9,10^.

The precise conceptual representation of variation is important for the application of computational methods in assessing human genomic variation in a clinical context. When studying rare diseases and cancers, clinical evaluation of patients increasingly includes interrogation of patient genomes for variants of potential clinical significance. Often, these assays will be highly targeted to query only those specific regions of interest, providing only partial information for clinical reporting. In some cases, observation of a variant allele is reported only as “heterozygous” (the presence of at least two different alleles at a genomic locus), “homozygous” (multiple copies of an allele at a locus with no other alleles), or “hemizygous” (an allele describing a locus for which there is only one total allele). These reports often omit further information regarding the total number of alleles at the locus or (for heterozygous variants) the composition of other alleles.

These abbreviated representations of human genotypes are imprecise, implying a diploid genotype when the patient may have aneuploidy caused by large-scale structural variation^11^ and/or meiotic nondisjunction^12^, typically resulting in abnormal phenotypes and disease. Heterozygous genotypes described in this way further connote the presence of a reference-agreement allele, though this too is not necessarily the case. To complicate the matter further, the manner in which variants are reported relies on an understood meaning of terms such as *allele, genotype*, and *haplotype*, which have similar but distinct meanings across different genomic communities and laboratories.

Clinical evaluation of genomic biomarkers also extends to drug response evidence, which can vary widely between individuals. In order to better understand how genetic information contributes to this variability, the pharmacogenomics (PGx) community collected evidence to gauge how genetic variants within a patient contribute to the overall responsiveness of a patient to different drugs^13^. Evidence from PGx knowledgebases can provide important information regarding drug toxicity and response within a patient, allowing for a more personalized treatment^14^.

One class of biomarkers describing PGx knowledge are “Star (*) Alleles”, which were first used to identify or denote alleles within the CYP gene family^15^. The results of PGx assays are often reported as diplotypes (pairs of haplotypes) due to the human genome being diploid^10^. The association of diplotypes and phenotypes enables the identification of pharmacogenetic interactions. For the assessment of PGx diplotypes, the most widely used nomenclature system for PGx alleles is the domain-specific “star” (*) system^16^. Due to the complex nature of PGx alleles and clinical assays, there continues to be ambiguity that can make it difficult to utilize PGx data in practice^17–24^. Some of these challenges were highlighted by the Centers for Disease Control and Prevention’s (CDC) Genetic Testing Reference Material (<u>GeT-RM</u>) Coordination Program test for clinical PGx genetic testing^25,26^. The results of this study demonstrate many inconsistencies due to a lack of a unified and standardized nomenclature system and different PGx designs. To help overcome the challenges regarding PGx data, the Clinical Pharmacogenetics Implementation Consortium was created to help educate and facilitate the use of PGx data in clinical settings^19,27–29^. Despite this, challenges remain in aligning PGx Star Alleles and other clinical biomarker domains^30^. Notably, there is a “*” representation that is called a spanning deletion in VCF, describing overlapping deletion Alleles at sites of other variants in a VCF file^31^.

To address the challenge of aligning the disparate genotype variation representations found in clinical reports, existing genomic variant exchange formats, and the PGx community, the Global Alliance for Genomics and Health (GA4GH)^32^ Genomic Knowledge Standards (GKS) Work Stream developed a Genotype model for the Variation Representation Specification (VRS; vrs.ga4gh.org)^33^ to enable the reliable and precise exchange of genotype variation between computer systems. The GA4GH VRS standard leverages a clearly defined terminology and information model, a value object design philosophy, and fully-specified JSON Schema, which allows it to meet these diverse use cases through modular variation representation. In this manuscript we describe this new model for representing genotypes using VRS, and demonstrate applications of this model to structure related concepts in other systems, including VCF, HGVS, and PGx Star Alleles.

## 2. Results

### 2.1. A landscape analysis of genotype concepts across communities

We first surveyed the requirements of genotype variation data as represented by large-scale genomic data standards (i.e. VCF), clinical reports (HGVS), and knowledgebases containing PGx (Star Allele) and/or variant-disease evidence (HGVS). We analyzed the conceptual alignment of terms from each specification to existing concepts in VRS to inform a conceptual framework for genotype representation (**Figure 1**).

**Fig 1.**
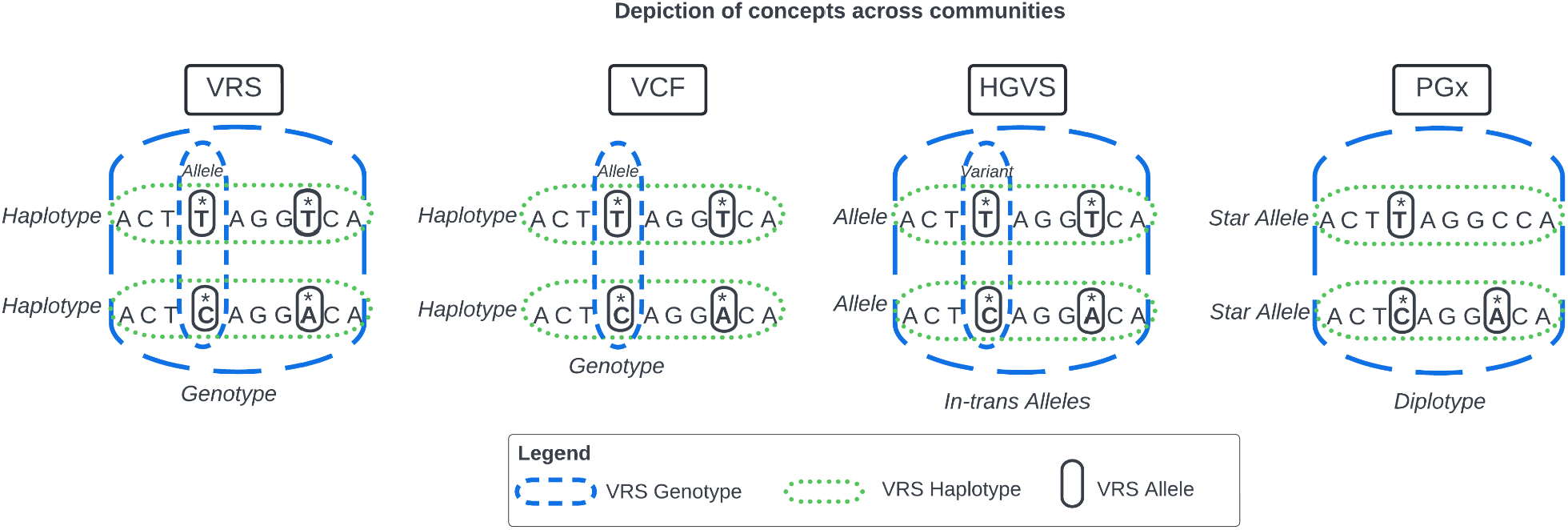
Genomics Concepts across Communities Communities use different terms for similar concepts. These concepts are represented with respect to VRS nomenclature while using terminology from each community. Among these standards, the VRS Genotype (blue dashes) aligns most closely to in-trans HGVS Alleles, VCF genotypes, and PGx diplotypes. Similarly, HGVS Alleles, VCF Haplotypes, and PGx Star Alleles are all aligned to the VRS Haplotype (green dots). Finally, a VRS Allele is conceptually aligned with a VCF Allele and an HGVS variant (black circles).

In VRS, an Allele is defined as the state of a molecule at a contiguous segment of a biological sequence. Variant Call Format (VCF) files store the variants identified by the sequencing platform and software used when analyzing a biological specimen. Each record within a VCF contains an identified variant with its corresponding position and the reference or wild type allele it was called against, along with other relevant information including the genotype. The VCF specification defines an allele as, “representing single genetic haplotypes (A, T, ATC)”, which aligns with the VRS definition of an Allele and the NCBI definition of a Contextual Allele^3^. The HGVS nomenclature uses the aligned term “variant” to describe a contiguous state on a sequence but uses the term “Allele” differently (see below). The PGx nomenclature also uses a broader definition of an allele (also discussed below).

A VRS Haplotype is defined as a set of non-overlapping Allele members that co-occur on the same molecule (they are in cis). A haplotype in the VCF specification is, “a set of variants which are known to be on the same chromosome in the germline genome”. This aligns to the VRS haplotype concept (defined as a set of Alleles “occurring on the same physical molecule”) and the ClinGen concept of a “haplotype”. The HGVS definition of an “allele” is “a series of variants on one chromosome.” An HGVS allele may represent a series of changes on the same physical sequence (i.e., in *cis*, akin to a VRS Haplotype), and a set of changes on different physical sequences (i.e., in *trans*, akin to a VRS Genotype). In addition, an HGVS allele may represent a set of changes with uncertain phase.The PGx community also uses the term “allele” to represent a concept analogous to VRS Haplotypes.

To model a Genotype in VRS, we built upon these concepts and analyzed the use of “genotype” or similar terms as described in other community standards. The official definition of a genotype as used in a VCF file is: “an assignment of alleles for each chromosome of a single named sample at a particular locus.” The reference allele in a VCF is encoded using a 0, while alternate alleles use 1, 2, etc. For example, in a diploid variant call, a heterozygous reference and alternate allele genotype would be encoded as 0/1 or a heterozygous alternate 1 and alternate 2 allele genotype would be encoded as 1/2. A homozygous alternate allele genotype is annotated as 1/1. Haploid variant calls only contain a single allele, while a triploid variant call would contain three alleles (e.g 0/0/1). An unphased genotype is represented using the “/” whereas a genotype with known phasing uses a “|” (e.g. 1 | 0).

The HGVS nomenclature doesn’t use the term genotype, but a user can create in-trans alleles which are conceptually aligned with the common meaning of the term (hgvs.org/DNA/variant/alleles). The use of “heterozygous” and “homozygous” as free text are used in some clinical reports^34^ accompanying an HGVS variant, in lieu of a formal HGVS trans-allele structure. This observation illuminated a key modeling requirement to capture the concept of heterozygous alleles within a genotype while lacking complete information about the constituent members.

We evaluated how PGx Star Alleles were represented within genotypes, and found that PGx evidence may be associated with a specific genotype representation described as a diplotype (a diploid genotype). Similarly, PGx evidence at the Star Allele level can be described naturally by a VRS Haplotype. This conceptual design benefits from a diploid constraint, and was well-suited to our starting model for Genotype (see **Methods**). We kept these diplotypes as an example case for testing in developing a VRS Genotype model.

### 2.2. The VRS Genotype information model and supporting classes

To develop the Genotype information model in VRS, we evaluated the definitions and constraints of the Allele and Haplotype models identified in our landscape analysis. The VRS Haplotype class had previously been defined as “a set of non-overlapping Allele members that co-occur on the same molecule”, but Haplotypes were allowed to contain a minimum of one Allele, designed to capture a semantic distinction between an Allele and a single-Allele Haplotype. However, after evaluating related concepts in the community, it was decided that the Haplotype information model should be updated to require at least two Allele members. This was informed by the lack of a distinction between a single-Allele Haplotype and an isolated Allele in other systems.

As a result of our modeling, we defined Genotype as “a quantified set of in-trans Molecular Variation at a genomic locus”, where *Molecular Variation* collectively refers to VRS Alleles, Haplotypes, and future classes of variation that exist on a contiguous molecule. This is in contrast to VRS *Systemic Variation* (including concepts such as Genotype and Copy Number Variation) which describe variation across several molecules within a system. We aligned this genotype definition with an information model that is flexible enough to capture the cross-domain concerns identified in our landscape analysis.

Each Molecular Variant constituting a Genotype is contained within an associated *Genotype Member* object to quantify the Molecular Variant present at a genomic locus (**Figure 2**). This provides a convenient mechanism for compactly representing identical Molecular Variation at a locus as well as expressing uncertainty in the count of that variation through the application of Definite Range or Indefinite Range objects. The count attributes of the Genotype Member and Genotype classes also enable compact representation of Molecular Variation in polyploid genomes and reflect similar conceptual structures designed for this purpose^35^.

**Fig 2.**
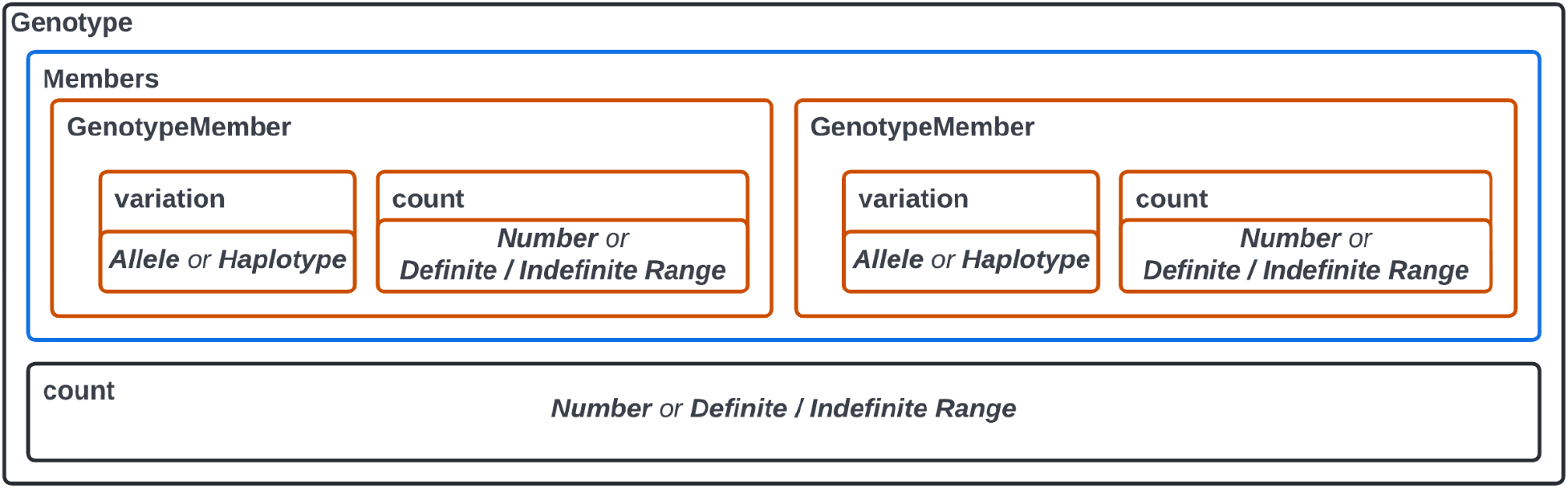
Genotype Class in VRS The Genotype class in VRS must contain at least one member consisting of an allele or haplotype and its count of occurrences within the system. This can be represented by an integer or as a definite/indefinite range. The Genotype also has a count, representing the expected total of the genotype’s molecule in the system, expressed as an integer or as a definite/indefinite range. This allows the user to describe what is known regarding the genotype without making an inference. For example, a user could add a single Genotype Member with a count = 1 and have the Genotype count = 2 to represent that there are additional molecular variations expected to exist but they are not explicitly indicated by the user or data.

In addition, a count field exists at the Genotype level for expressing the total copies of the genomic locus as described by the Genotype Members. The Genotype count value could be greater (but never less) than the summation of counts across Genotype Members. In such cases, the difference conveys additional unspecified Molecular Variation that is expected to exist but is not explicitly represented. This feature allows for precisely representing ambiguity in genotype concepts when not all Molecular Variation are reported.

### 2.3. Applications of the Genotype information model

We evaluated how this structure provides the flexibility to represent concepts from a simple two allele genotype or a diplotype composed of a single Allele in-trans with a haplotype. The two-Allele genotype example is exemplified by a common VCF record pattern, where two or more VCF Alleles are expressed in-trans independent of in-cis phasing with neighboring Alleles (e.g. 0/1). In this case, each VCF Allele is expressed as a VRS Allele, put into a Genotype Member object with count=1, and both of those Genotype Members added to a Genotype with count=2 (**Figure 3A**). We also developed a utility for annotating VCF records with VRS Alleles (see **Methods**) to assist Genotype reconstruction from single-sample and multi-sample VCFs.

**Fig 3.**
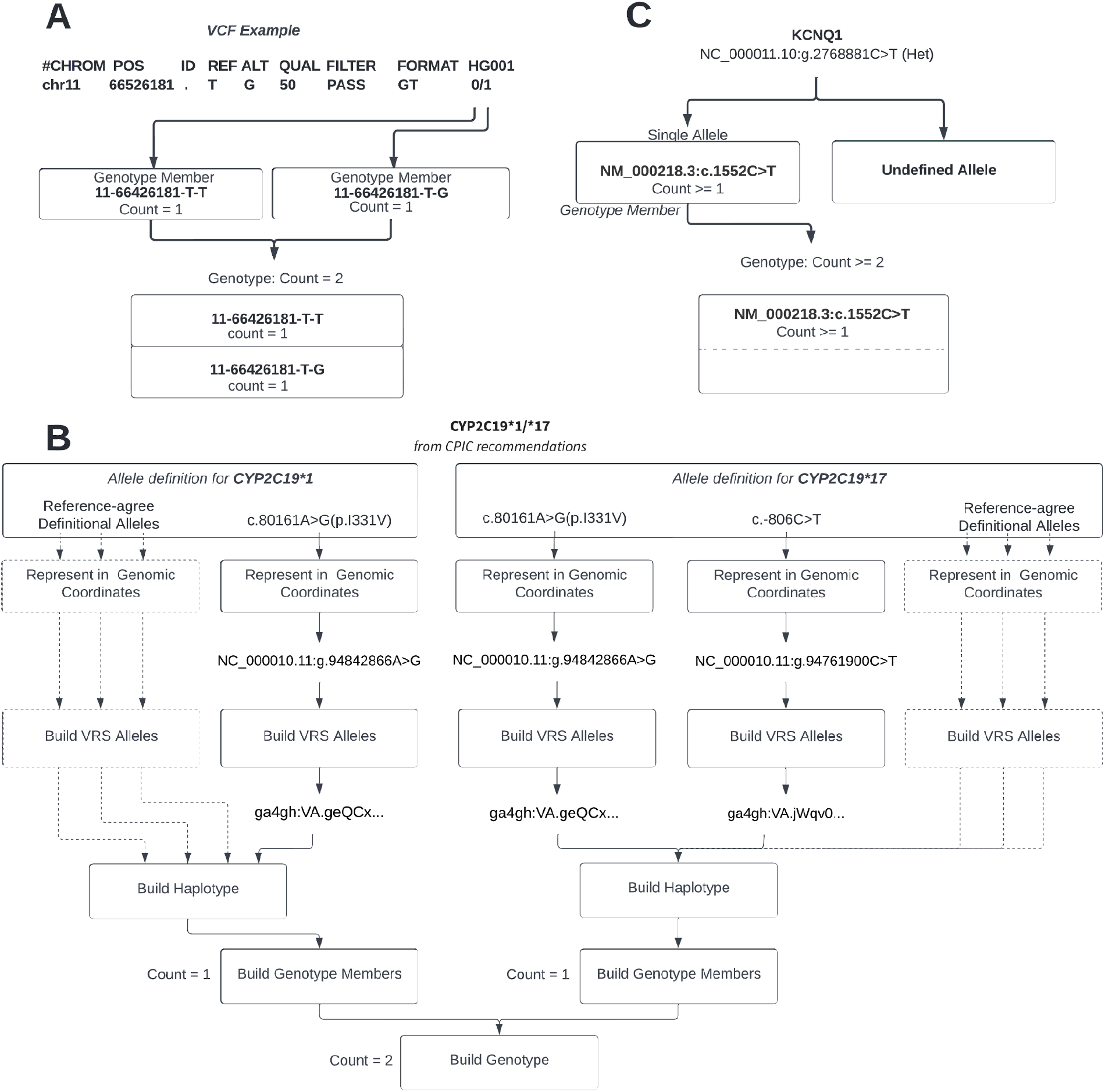
Visualization of Genotypes in VRS Variants are represented in their genomic coordinates and then normalized and translated into their VRS-allele ID’s using VRS-Python. **A**. Representation of a 0/1 Genotype from a VCF. **B**. CYP2C19*1 is composed of a single variant and can be immediately placed into a Genotype Member with a copy state of 1. CYP2C19*17 contains two variants in-cis which needs to be represented by a Haplotpye and then placed into a Genotype Member with count = 1. These two genotype members are then used to construct the genotype shown above with a total copy count of 2. A Star Allele representation incorporating reference-agree VRS Alleles is depicted with dashed lines. **C**. Representation of a heterozygous variant from an eMerge report.

A more complex scenario was tested on the *CYP2C19 *1/*17* diplotype (**Figure 3B**) as represented by changes from a reference sequence. Initialization of this process requires selection of a sequence context for describing the constituent variants. In this example we selected the GRCh38 genomic reference^36^. It is important that a genomic DNA sequence is used in this step, as Star Alleles include variation in regulatory and intronic regions and representation of intronic variation with respect to a cDNA sequence (e.g. RefSeq NM_ sequences) is dependent upon an inferred alignment of these variants to a genomic reference. VRS Alleles were constructed on the selected reference sequence, and in-cis Alleles were subsequently grouped into VRS Haplotypes. The count of each Molecular Variation (in this example, one *<u>CYP2C19 *1</u>* Star Allele and one *<u>CYP2C19 *17</u>* Star Allele) is specified using the Genotype Member class. These Genotype Members are assembled into a Genotype and the overall count (2) of alleles at the locus is recorded, explicitly indicating a diploid state at this locus.

Nuances to the use and meaning of the VRS Genotype model for representing Star Alleles were captured in discussion with members of the PGx informatics community. While the VRS Genotype model faithfully represents the variants for these Star Alleles as displayed in PharmVar, the meaning of these PGx Star Alleles and how they should be assessed is more complex than simply observing the described collection of non-reference variants. The Star Allele model also assumes that there is an associated set of definitive locations that have been assayed (and are expected to be reference-agree) to properly assign Star Allele Haplotypes from patient sequencing data. To address this, we leveraged the Allele design of VRS to demonstrate a data structure to efficiently communicate this nuance between systems using both variant and reference-agree Alleles (**Figure 3B** and **Methods**). This has the added benefit of preserving the context under which Star Alleles are described, aiding reinterpretation and data reuse as additional Star Alleles are discovered and the number of definitive sites increase.

Finally, we tested this model on Genotypes with missing members to illustrate how this model captures those annotations. Starting with an eMERGE-seq panel report^34^, we create a Genotype from a heterozygous variant report with only one allele described. We used the VRS Indefinite Range concept to express the heterozygous variant as observed at least once at a genomic locus with at least two alleles (**Figure 3C**). An alternative could also be to infer a diploid state for this report, in which case we would represent this as a variant observed once at a locus of two alleles.

### 2.4. Implementation support

The definition and information model for Genotypes has been implemented in documentation at vrs.ga4gh.org/en/genotype-draft/terms_and_model.html#genotype, structured in JSON Schema at github.com/ga4gh/vrs/blob/genotype-draft/schema/vrs.json and implemented in Python at github.com/ga4gh/vrs-python/tree/pgx. We have also created example PGx jupyter notebooks to demonstrate how to create and use Genotypes and other VRS components within VRS-Python to build and search Star Alleles at github.com/ga4gh/vrs-python/blob/pgx/notebooks/PGx.ipynb.

In addition to the static examples available above, this and other VRS-Python notebooks can be run from a local copy of the vrs-python repository or using zero-install cloud-based notebooks hosted at mybinder.org/v2/gh/ga4gh/vrs-python/pgx. The cloud-based notebooks are a simple mechanism for newcomers to interactively test the functionality and scope of VRS-Python and associated VRS models by leveraging our publicly accessible REST APIs to support services. A user may follow the examples provided within the notebooks to gain an understanding of VRS and can even edit or add cells to further explore VRS using their own data or examples.

## 3. Discussion

Defining a model for genotype representation required careful conceptual alignment and semantic precision for interoperability of this model with similar concepts across different communities. We found that while the VRS, VCF, HGVS, and PGx communities have some differences between the terms allele, haplotype, and genotype, there are shared conceptual relationships describing the in-cis and in-trans representation of sequence variants at a genomic locus. We found that these shared conceptual models enabled a unified computational structure for interchangeable and lossless description of these concepts between systems, advancing our ability to automate scalable evidence search operations between assayed data and genomic knowledgebases.

The VRS Genotype model explicitly captures the count of individual alleles and all expected alleles at a locus as independent values, allowing for the flexible description of genomic loci and enabling precise forms of ambiguity using VRS Definite Range and Indefinite Range quantifiers. We demonstrated how this allows for reconstruction of ambiguity as derived from clinical reports and representation of Genotypes of ambiguous ploidy. We also illustrated how this model enables lossless capture of the VCF record-level genotype model, and like VCF this provides a straightforward mechanism for representation of alleles at polyploid loci. In addition, we showed how the Genotype model enables the representation of variants as expressed in PharmVar. We also illustrated how this model can be extended using the modular design of VRS to associate Genotypes with additional necessary elements for defining PGx Star Alleles. Together, these findings provide a template for the flexible use of VRS Genotypes across various genomics communities with domain-specific requirements.

There is an ongoing need for the standardization of information models and schema for assay metadata such as “assayed regions”. We demonstrated a method to address some of these needs by leveraging the modularity of VRS to construct a computable genotype with a Star Allele nomenclature label. We will also be improving our VCF-annotation tool to include the ability to annotate VRS genotypes in VCF files and will be applying the VRS genotype model to the ClinVar database. The VRS Genotype model is a completed draft model under community review and is anticipated to be finalized by September 2022 and included in the next official minor version release of VRS.

Prior to this work, data exchange between the pharmacogenomics and other genomic communities has been somewhat challenging. VRS allows us to precisely describe the genotypes within PGx data, VCF files, and lab reports. Therefore, opening an avenue for advanced queries and search operations from many different pharmacogenomic and clinical assays and knowledgebases through the use of federated data.

## 4. Methods

### 4.1. Community modeling and use case discussions

The Genotype model was initially discussed and revisited on several occasions during the development of VRS, and an initial model was under consideration for the VRS 1.2 release. This initial model was a structure containing a set of Haplotypes and was designed to represent the set as an *in-trans* model. This model was unwieldy due to the lack of support for Molecular Variation counts or total Molecular Variant count at a locus.

In July 2022, the GA4GH sponsored a VRS hackathon at the Intelligent Systems for Molecular Biology 2022 Annual Conference in Madison, Wisconsin. During the hackathon, modeling of the Genotype class was selected as a preferred topic, and participants in this activity worked together to evaluate the Genotype model and its relation to similar concepts in different communities, including immunogenomic and pharmacogenomic use cases. The group discussed the concepts of alleles, genotypes, and haplotypes and how they are related to one another to determine the best way to precisely model a genotype within VRS. Multiple examples from clinical reports, genomic assay results, and genomic knowledgebases were chosen to test and revise the ideas proposed. Once the group finalized the VRS Genotype model, they used the model to describe PGx alleles using VRS to test the model for interoperability between assayed PGx data and pharmacogenomic knowledge bases.

### 4.2. Community Review

Community involvement and review is a critical component of developing standards that are meant for the global community. We presented the new VRS genotype model during the July 18th and July 25th GA4GH Variation Representation meetings, and with the VCF community maintainers on the GA4GH July 27th VRS/VCF alignment call to receive feedback from interested community members and domain experts. We also sent an open call for review to the GA4GH community for comments and questions during our open review period. The process of responding to community feedback is expected to be completed by September 2022. The community comments for the review of this model were documented online at github.com/ga4gh/vrs/pull/394.

### 4.3. VRS-VCF annotation tool

The VRS-VCF annotation tool allows users to annotate the reference and alternate alleles of a VCF record with VRS. The VRS allele identifier is stored in the INFO field of the VCF and an optional pickle file containing the entire VRS object can be created for all the annotated records. The VRS allele identifier can then be used for precise and speedy lookup of information from databases utilizing VRS, which drastically simplifies the variant annotation process. The tool is open-source and readily available online at github.com/ga4gh/vrs-python/blob/main/src/ga4gh/vrs/extras/vcf_annotation.py.

### 4.4. Software availability

All code supporting the development, documentation, implementation, and validation of the VRS Genotype model is available online at <u>GitHub</u>as indicated throughout the text, under the permissive Apache 2.0 open source license.

## 5. Acknowledgments

The authors thank Li Gong, Teri E. Klein, Ryan Whaley, and Michelle Whirl-Carillo (Stanford University) for important discussions and critical feedback that substantially advanced this work. WAG & AHW were supported by the National Human Genome Research Institute (NHGRI) award R35HG011949. LB & HR were supported by the NHGRI award U24HG006834. MSC was supported by the National Cancer Institute (NCI) award U01CA242954-01. RRF & KR were supported by the NHGRI award U41HG006834. RRF was supported by the NHGRI award R35HG011899. KR was supported by the NHGRI awards U41HG009649, U41HG009650. ADY was supported by the Wellcome Trust [WT222155/Z/20/Z] and the European Molecular Biology Laboratory. RKH was supported by ClinGen, Invitae, Inc, and MyOme, Inc..

For the purpose of open access, the author has applied a CC-BY public copyright licence to any author accepted manuscript version arising from this submission.

## Notes

### Competing Interest Statement

The authors have declared no competing interest.

## Bibliography

1. Danecek, P. et al. The variant call format and VCFtools. Bioinformatics 27, 2156–2158 (2011).

2. den Dunnen, J. T. et al. HGVS Recommendations for the Description of Sequence Variants: 2016 Update. Hum. Mutat. 37, 564–569 (2016).

3. Holmes, J. B., Moyer, E., Phan, L., Maglott, D. & Kattman, B. SPDI: data model for variants and applications at NCBI. Bioinformatics 36, 1902–1907 (2020).

4. Pawliczek, P. et al. ClinGen Allele Registry links information about genetic variants. Hum. Mutat. 39, 1690–1701 (2018).

5. Sherry, S. T. et al. dbSNP: the NCBI database of genetic variation. Nucleic Acids Res. 29, 308–311 (2001).

6. Fokkema, I. F. A. C. et al. LOVD v.2.0: the next generation in gene variant databases. Hum. Mutat. 32, 557–563 (2011).

7. International Standing Committee on Human Cytogenomic Nomenclature. ISCN 2020: An International System for Human Cytogenomic Nomenclature (2020). (Karger, 2020).

8. Quinlan, A. R. & Hall, I. M. BEDTools: a flexible suite of utilities for comparing genomic features. Bioinformatics 26, 841–842 (2010).

9. Wagner, A. H. et al. A harmonized meta-knowledgebase of clinical interpretations of somatic genomic variants in cancer. Nat. Genet. 52, 448–457 (2020).

10. Kalman, L. V. et al. Pharmacogenetic allele nomenclature: International workgroup recommendations for test result reporting. Clin. Pharmacol. Ther. 99, 172–185 (2016).

11. Escaramís, G., Docampo, E. & Rabionet, R. A decade of structural variants: description, history and methods to detect structural variation. Brief. Funct. Genomics 14, 305–314 (2015).

12. Angell, R. First-meiotic-division nondisjunction in human oocytes. Am. J. Hum. Genet. 61, 23–32 (1997).

13. Jones, D. S. How personalized medicine became genetic, and racial: Werner Kalow and the formations of pharmacogenetics. J. Hist. Med. Allied Sci. 68, 1–48 (2013).

14. Sim, S. C., Altman, R. B. & Ingelman-Sundberg, M. Databases in the area of pharmacogenetics. Hum. Mutat. 32, 526–531 (2011).

15. Nebert, D. W. Suggestions for the nomenclature of human alleles: relevance to ecogenetics, pharmacogenetics and molecular epidemiology. Pharmacogenetics 10, 279–290 (2000).

16. Robarge, J. D., Li, L., Desta, Z., Nguyen, A. & Flockhart, D. A. The Star-Allele Nomenclature: Retooling for Translational Genomics. Clinical Pharmacology & Therapeutics vol. 82 244–248 (2007).

17. Swen, J. J. et al. Translating pharmacogenomics: challenges on the road to the clinic. PLoS Med. 4, e209 (2007).

18. Scott, S., Abul-Husn, N., Obeng, A. O., Sanderson, S. & Gottesman, O. Implementation and utilization of genetic testing in personalized medicine. Pharmacogenomics and Personalized Medicine 227 (2014) doi:10.2147/pgpm.s48887.

19. Relling, M. V. & Klein, T. E. CPIC: Clinical Pharmacogenetics Implementation Consortium of the Pharmacogenomics Research Network. Clin. Pharmacol. Ther. 89, 464–467 (2011).

20. Lee, K. C., Ma, J. D. & Kuo, G. M. Pharmacogenomics: Bridging the gap between science and practice. Journal of the American Pharmacists Association vol. 50 e1–e17 (2010).

21. Ma, J. D., Lee, K. C. & Kuo, G. M. Clinical application of pharmacogenomics. J. Pharm. Pract. 25, 417–427 (2012).

22. Stanek, E. J. et al. Adoption of pharmacogenomic testing by US physicians: results of a nationwide survey. Clin. Pharmacol. Ther. 91, 450–458 (2012).

23. Malentacchi, F. et al. Is laboratory medicine ready for the era of personalized medicine? A survey addressed to laboratory directors of hospitals/academic schools of medicine in Europe. Clinical Chemistry and Laboratory Medicine (CCLM) vol. 53 (2015).

24. Hess, G. P., Fonseca, E., Scott, R. & Fagerness, J. Pharmacogenomic and pharmacogenetic-guided therapy as a tool in precision medicine: current state and factors impacting acceptance by stakeholders. Genet. Res. 97, e13 (2015).

25. Pratt, Zehnbauer, Wilson & Baak. Characterization of 107 genomic DNA reference materials for CYP2D6, CYP2C19, CYP2C9, VKORC1, and UGT1A1: a GeT-RM and Association for …. The Journal of molecular.

26. Pratt, V. M. et al. Characterization of 137 Genomic DNA Reference Materials for 28 Pharmacogenetic Genes. The Journal of Molecular Diagnostics vol. 18 109–123 (2016).

27. Gaedigk, A., Whirl-Carrillo, M., Pratt, V. M., Miller, N. A. & Klein, T. E. PharmVar and the Landscape of Pharmacogenetic Resources. Clin. Pharmacol. Ther. 107, 43–46 (2020).

28. Whirl-Carrillo, M. et al. Pharmacogenomics knowledge for personalized medicine. Clin. Pharmacol. Ther. 92, 414–417 (2012).

29. Relling, M. V. et al. New Pharmacogenomics Research Network: An Open Community Catalyzing Research and Translation in Precision Medicine. Clin. Pharmacol. Ther. 102, 897–902 (2017).

30. Caudle, K. E. et al. Standardization can accelerate the adoption of pharmacogenomics: current status and the path forward. Pharmacogenomics 19, 847–860 (2018).

31. GATK Documentation Team. Spanning or overlapping deletions (* allele). Genome Analysis Toolkit https://gatk.broadinstitute.org/hc/en-us/articles/360035531912-Spanning-or-overlapping-deletions-allele-.

32. Rehm, H. L. et al. GA4GH: International policies and standards for data sharing across genomic research and healthcare. Cell Genom 1, 100029 (2021).

33. Wagner, A. H. et al. The GA4GH Variation Representation Specification: A computational framework for variation representation and federated identification. Cell Genomics 1, (2021).

34. Murugan, M. et al. Genomic considerations for FHIR®; eMERGE implementation lessons. J. Biomed. Inform. 118, 103795 (2021).

35. Clevenger, J. P., Korani, W., Ozias-Akins, P. & Jackson, S. Haplotype-Based Genotyping in Polyploids. Front. Plant Sci. 9, 564 (2018).

36. Schneider, V. A. et al. Evaluation of GRCh38 and de novo haploid genome assemblies demonstrates the enduring quality of the reference assembly. Genome Res. 27, 849–864 (2017).

